# FMAGJ: Fluorescence Microscopic Astrocyte Gap Junction Images Dataset

**DOI:** 10.64898/2025.12.25.695956

**Authors:** Evgeniya Kirichenko, Rozaliia Nabiullina, Ekaterina Churkina, Konstantin Avakyan, Julia Gordeeva, Darya Sedova, Chizaram Nwosu, Sergey Golovin, Maria Kaplya, Alina Sereda, Stanislav Rodkin, Mikhail Petrushan

## Abstract

This work introduces a novel and specialized dataset of high-resolution fluorescence microscopic images focused on astrocytic gap junctions, aiming to get insights into intercellular communication in both healthy and pathological brain conditions. The dataset includes 20 z-stack image series—10 from human glioblastoma tissue and 10 from healthy rat brain tissue—each containing between 22 and 104 optical slices. These images were acquired using standardized protocols following immunofluorescence labeling with antibodies against connexin 43 (Cx43), enabling visualization of gap junction localization at the subcellular level.

The dataset is tailored to support detailed morphological and quantitative analyses of gap junction networks, featuring metadata including species and localization of possible gap junctions in images provided by bounding boxes and image masks. Its structure facilitates comparative studies across physiological states and species, enhancing translational and evolutionary perspectives on astrocytic connectivity. Given the labor-intensive nature of manual gap junction quantification, this dataset serves as a resource for the development of machine learning tools capable of automating the detection and analysis of Cx43-positive signals.

**CCS CONCEPTS:** Computing methologies∼Computer vision problems∼Object detection • Applied computing∼Life and medical sciences∼Computational biology/Molecular structural biology • Software and its engineering∼Software notations and tools∼Software libraries and repositories

## 1 INTRODUCTION

Brain astrocytes play a fundamental role in maintaining neuronal homeostasis, regulation of synaptic conduction, and neuronal plasticity through the constant release and uptake of neurotransmitters. An essential component to enable these processes is intercellular communication. Through the formation of intercellular gap junctions, the astroglia acquires the function of a spatial buffer for the regulation of extracellular ionic and metabolite concentration and homeostasis, as well as the regulation of soma volumes of the astrocyte itself and its outgrowths [1]. Gap junctions are represented by a system of tightly packed channels penetrating the bilipid layers of the membranes of two contacting cells. Each channel consists of two docked half-channels called connexons, and the subunit of each is the protein connexin. In the mammalian brain, astrocytic gap junctions are composed primarily of connexin 43(Cx43), less frequently connexin 30. As a monomer, Cx43 consists of four hydrophobic transmembrane domains, two extracellular loops, one cytoplasmic loop, and two N-C free end domains of the molecule. Astrocytes connected via Cx43 containing gap junctions form a kind of functional glial syncytium or gap junction-mediated glial networks [2]. Degradation of such functional networks is associated with a number of neurodegenerative diseases such as Alzheimer’s disease, Parkinson’s disease, brain atherosclerosis, stroke, multiple sclerosis, demyelinating diseases, epilepsy, prion diseases and various psychiatric syndromes [3]. Separately, there is a relation of Cx43 consisting gap junctions with the growth of glial brain tumors, including fatal glioblastoma [4]. The first works in this area suggest the formation of a new view on the heterogeneity of Cx43 expression in brain tumors and the related promising new opportunities in the diagnosis and treatment of CNS neoplasms [5]. At the same time, the role of gap junctions in the formation of functional networks and cell ensembles, as well as their involvement in information processing in the normal pathology-free brain still remains unclear. Understanding of gap junctional role as well as the role of Cx43 protein will be facilitated by studies of detection of Cx43 positive signals bound to astrocyte compartments throughout the volume in various structures of the normal pathology-free brain as well as in tumor-affected brain tissues. Such studies are expedient for the development of tools for the search and creation of new therapeutic strategies based on the regulation of connexin expression and the activity of gap junctions and half-channels in human brain tumors. Thus, fast and efficient detection of minor changes in systems of gap junction is not possible without specialized machine learning technologies. To implement such specialized neural network solutions, it is necessary to collect an image dataset consisting of multiple Z-stack of astrocytes and Cx43 positive signals in both glioma tumor tissue and normal pathology-free tissue. Sample preparation involved immunofluorescence with antibodies against Cx43 proteins. In this case, to investigate the characterization of the gap junction system without tumor transformation, healthy control samples were obtained from adult laboratory rats with absence of gliomas.

This work presents a unique dataset of fluorescence microscopic images focused specifically on astrocyte Cx43 puncta. Our dataset addresses a significant gap in the field of neuroscience research by providing high-resolution, specialized images of Cx43 puncta, which are crucial for understanding gap junction astrocytic communication networks in the brain. (see APPENDICES A.1)

### 2 RELATED DATASETS

A systematic review of publicly available astrocyte image datasets reveals that while several collections exist, they generally focus on broader aspects of astrocyte morphology. These datasets differ from ours in terms of imaging modalities, resolution, staining protocols, and biological focus, rendering them unsuitable for direct mixing with our data for Cx43 expression analysis purposes (e.g. for training machine learning models on mixed data). (see APPENDICES A.2)

#### 2.1 Key features of Our Dataset

Our dataset provides high-resolution images of individual astrocytes with clear visualization of gap junctions and key features:

1. Specialized Focus on Gap Junctions: Unlike existing datasets that emphasize whole-cell astrocyte morphology or general detection, our collection is uniquely tailored to visualize and analyze intercellular Cx43 puncta.

2. High-Resolution Single-Cell Imaging: The dataset includes high-magnification images of individual astrocytes, enabling subcellular-level morphological studies. This contrasts with other datasets, such as the Annotated Astrocyte Dataset [7], which often contain multiple cells per image.

3. Pathology-Control Comparisons: Samples are derived from both normal brain tissue and pathological conditions, particularly human glioblastoma, enabling comparative studies relevant to disease mechanisms.

4. Despite the seemingly small size of the subdomains, the number of images is greater than all the available connexin mask layouts.

The comparative assessment confirms that no currently available dataset [7], [8], [9], [10] offers the same combination of specificity, resolution, and metadata richness for studying astrocyte gap junctions. As such, this dataset represents a valuable addition to the field, enabling deeper exploration of astrocytic communication in both physiological and pathological contexts. The comparison of datasets is shown in APPENDIX A2.

### 3 DATA COLLECTION

#### 3.1 Sample Preparation Protocol for Normal Brain Tissue

S Q Qections of rat brain tissue were used as an object of study. Animals weighing 150–180 g and age 0.5 years were used. The animals’ brains were fixed by transcardial perfusion method. After weighing, the animals were anesthetized with 5 mg/kg xylazine and 50–100 mg/kg zoletil intramuscularly and secured on the perfusion table with adhesive tape. After venous injection of 0.1 ml of heparin, an operation to open the chest was performed, the lower parts of the right and left ventricles were cut off, freeing their lumen. A cannula connected to a syringe with a perfusion solution was inserted into the aorta through the left ventricle and was fixed there. Perfusion was performed firstly with a phosphate buffer solution, pH 7.4, brought to 37 °C (Sigma-Aldrich, USA), and then with a cooled fixative solution of 4% paraformaldehyde (Sigma, EMS, USA) in phosphate buffer (pH 7.4). The perfusion rate corresponded to the average speed of blood movement through the vessels; the total volume of perfused fluids averaged 400 ml. After the end of perfusion, the animal was left for 2 hours at room temperature, then, the brain was removed and placed in 4% paraformaldehyde for post-fixation overnight at a temperature of 4 °C.

In the rat brain, the objects of study were the areas of the cortical (barrel cortex S1) and thalamic (nuclei VPM, VPL, RT) representation of the vibrissae, as well as the hippocampus. After postfixation, a section was isolated from the brain according to the coordinates: the first incision was 0.2 mm rostral from bregma, the second incision was 6.04 mm caudal from bregma, while the brain was not dissected laterally. The excised brain fragment was glued with the caudal side of the slice down to the stage of a VT 1000E vibratome (Leica, Germany). Next, frontal 40 μm sections were prepared. Every section was placed in a drop of phosphate buffer in a Petri dish and viewed under a stereotactic loupe. As soon as the objects under study were identified on the frontal sections in accordance with the atlas (barrel cortex, thalamic nuclei, hippocampus), cutting stopped.

For cryoprotection, sections were placed in a solution of 15% sucrose for 10 minutes, then 30% sucrose for 10 minutes. Unmasking of antigenic activity was carried out by instant freezing of sections on foil at -80 °C and subsequent thawing in drops of phosphate buffer. This process was followed by washing the sections in three changes of fresh phosphate buffer (pH 7.4). Sections were incubated in a mixture of primary antibodies to glial fibrillar acidic protein (mouse monoclonal Glial fibrillar Acidic Protein, Sigma-Aldrich, USA, SAB5201104) at a dilution of 1/200 and to Cx43 protein (rabbit polyclonal, Elabscience, USA, E-AB-70097) diluted 1/200 for 7 days at 20 °C with the addition of 0.1% Triton-X and 0.1% sodium azide. Incubation was followed by washing the sections in three changes of fresh phosphate buffer (pH 7.4). Next, the sections were incubated for 24 hours at 10 °C in a mixture of secondary antibodies conjugated with a fluorescent label: 1) goat anti-rabbit conjugated with Abberior STAR ORANGE (Abberior, Germany), diluted 1/200; 2) goat anti-mouse conjugated with Abberior STAR RED (Abberior, Germany), diluted 1/200; and 3) with green dye Sytox green (Thermo Fisher Scientific, USA), diluted 1/1000. Finally, the sections were placed on a glass slide in antifade solution (Abberior, Germany) and covered with a coverslip, 0.17 mm of thickness. The edges of the coverslip were carefully packed with clear nail polish around the entire perimeter. Visualization of the obtained data was carried out using a high-resolution system (Abberior Facility Line, Abberior Instruments GmbH, Germany) in confocal laser scanning mode.

Due to ethical constraints precluding the acquisition of healthy human brain tissue, we utilized exclusively glioblastoma specimens for the pathological group. For comparative analysis, healthy animal samples were obtained from model rat organisms given that the structural and functional organization of Cx43 exhibits no fundamental interspecies differences between humans and rats, thereby justifying the use of rat tissue as an appropriate control.

#### 3.2 Sample Preparation Protocol for Brain Tumor Tissue

All tumor samples were obtained with a collaboration of neurosurgery department of the local medical university. Each sample was accompanied by written agreement from the patient or relative, the study was approved by the Bioethics of the university and accompanied by a pathomorphological diagnosis. For confocal microscopy, tumor samples measuring 3/3 mm were immediately placed during surgery in 4% paraformaldehyde by immersion fixation method (Sigma, EMS, USA) in phosphate buffer (pH 7.4). After 24 hours’ fixation, the tumor pieces were filled with a warm solution of 2.5% agarose gel; after cooling and hardening of the gel, square sections with enclosed tissue fragments were selected with a blade and glued to the vibratome table. Next, vibratome 40 µm sections were made on a VT 1000E vibratome (Leica, Germany).

For cryoprotection, sections were placed in a solution of 15% sucrose for 10 minutes, then 30% sucrose for 10 minutes. Unmasking of antigenic activity was carried out by instant freezing of sections on foil at -80 °C and subsequent thawing in drops of phosphate buffer. This was followed by washing the sections in three changes of fresh phosphate buffer (pH 7.4). Sections were incubated in a mixture of primary antibodies to glial fibrillar acidic protein (mouse monoclonal Glial fibrillar Acidic Protein, Sigma-Aldrich, USA, SAB5201104) at a dilution of 1/200 and to Cx43 protein (rabbit polyclonal, Elabscience, USA, E-AB-70097) diluted 1/200 for 7 days at 20 °C with the addition of 0.1% Triton-X and 0.1% sodium azide. Incubation was followed by washing the sections in three changes of fresh phosphate buffer (pH 7.4). Next, the sections were incubated for 24 hours at 10 °C in a mixture of secondary antibodies conjugated with a fluorescent label: 1) goat anti-rabbit conjugated with Abberior STAR ORANGE (Abberior, Germany), diluted 1/200; 2) goat anti-mouse conjugated with Abberior STAR RED (Abberior, Germany), diluted 1/200; and 3) with green dye Sytox green (Thermo Fisher Scientific, USA), diluted 1/1000. Finally, the sections were placed on a glass slide in antifade solution (Abberior, Germany) and covered with a coverslip, 0.17 mm of thickness. The edges of the coverslip were carefully packed with clear nail polish around the entire perimeter. Visualization of the obtained data was carried out using a high-resolution system (Abberior Facility Line, Abberior Instruments GmbH, Germany) in confocal laser scanning mode.

#### 3.3 FMAGJ dataset

The dataset comprises two distinct subdatasets derived from human glioblastoma tissue and healthy rat brain samples, respectively. In total, the dataset includes 1,058 samples, composed of 541 from human patients diagnosed with glioblastoma cancer and 517 from healthy laboratory rats. Each subdataset contains 10 Z-stacks that are reconstructing a full three-dimensional picture of astrocytes, resulting in 20 complete cell reconstructions across the dataset. Each subdataset is assigned a unique identifier to ensure traceability and structural clarity. Astrocyte imaging was conducted using a fluorescently labeled Cx43 marker, chosen for its role in labeling gap junction proteins. Fluorescence emission was captured using the Star Orange dye, providing clear contrast for subcellular structures.

The annotation process was carried out manually by one domain PhD expert with academic backgrounds in neural biology. Annotation tasks were performed using a PULLAR proprietary laboratory’s software tool [14] designed specifically for labeling objects. The criteria for identifying Cx43 puncta were based on the presence of high-intensity, sharply localized fluorescent signal regions, which were consistently interpreted by experts as indicative of connexin-rich junctions (Fig.1). The annotation procedure was conducted in segmentation mode using the Brush tool for a single object class, “Connexin43”.

**Figure 1.**
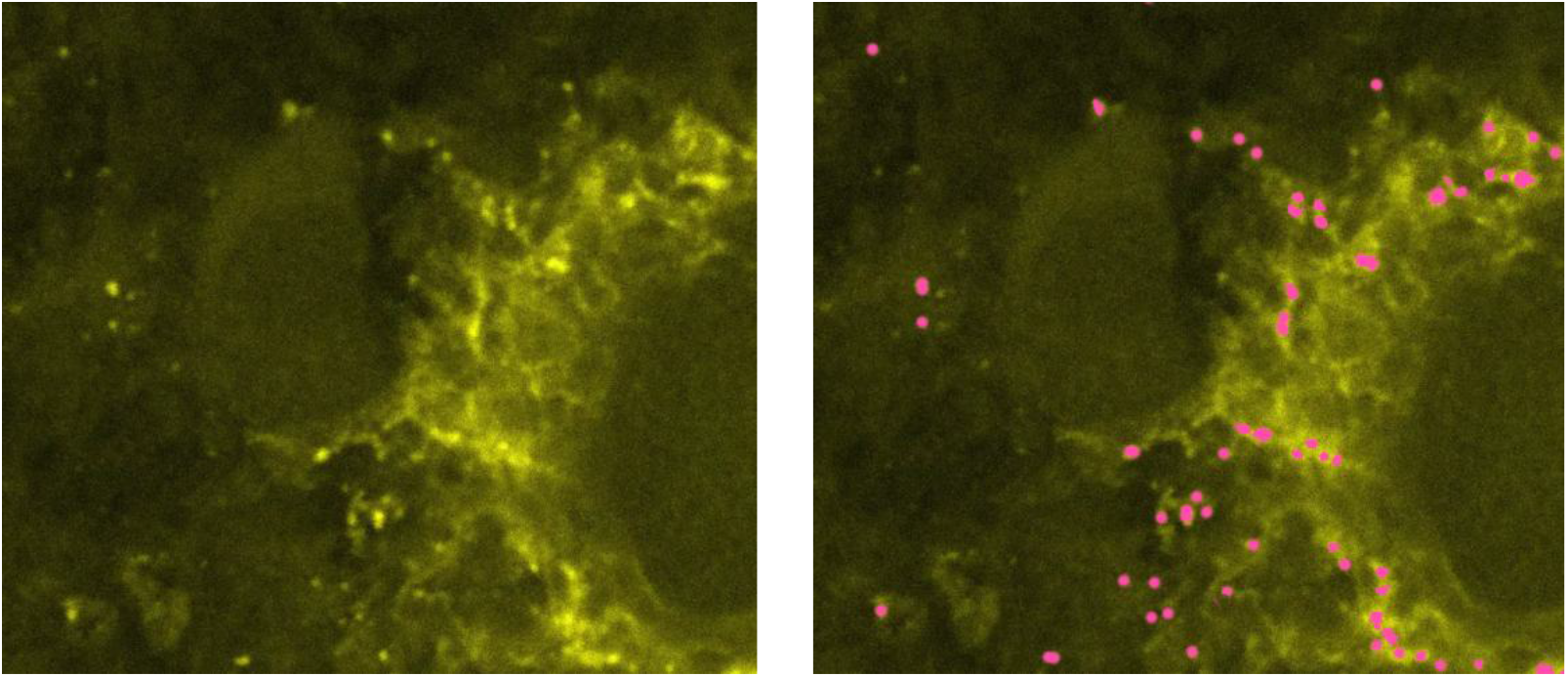
Example of ground truth mask labeling.

Binary object masks were obtained as a result of annotation. The conversion to COCO VIDEO format utilized OpenCV’s computer vision functions, which implement fundamental mathematical algorithms including: contour extraction based on Suzuki-Abe topological analysis, minimum bounding rectangle computation via coordinate extremum optimization, and connected component analysis using Union-Find data structures. This included geometric parameters of bounding boxes (bbox), object area, instance identifier (instance_id) for tracking, as well as frame and video sequence numbers. Matching of Cx43 puncta between consecutive slices was performed using the Hungarian algorithm for subsequent re-identification of individual object lifetimes throughout the entire Z-stack. The cost function incorporated two parameters: Intersection over Union (IoU) and Euclidean distance between bounding box centers. The threshold value for maximum distance was set at 10 pixels to exclude physically impossible object displacements between frames.

Data were structured as follows: annotations – Z-stacks with unique identifiers (UUID); frames – individual slices (images); tracks – Cx43 puncta also with shortened UUIDs. Each tracked object was assigned a unique identifier in the format [frame_number][image_prefix][4_hash_characters]. New objects appearing during observation received unique identifiers of the frame number at which observation of the corresponding object began.

Based on the obtained data, the following quantitative parameters were calculated: number of gap junctions per frame, observation duration of individual junctions, dynamics of Cx43 puncta formation and disappearance processes, and spatial distribution of objects in the microscope field of view.

Our datasets are posted at the following link: https://github.com/mdi-bbm/FMAGJ. The parameters are described in the documentation. (see APPENDICES A.3) The dataset is structured into two subdatasets: rat_healthy and human_glioblastoma. Each subdataset contains 10 unique Z-stack folder names. Within each folder, there is a single Z-stack scan consisting of: an images folder containing PNG images, a masks folder with initial binary segmentation PNG masks, and a coco_video_detection folder with generated detections in COCO format. The folder contains two JSON files: annotation.json with the bounding box annotations, and track_info.json with information on the duration of each Cx43 puncta across each Z-stack.

### 4 STATISTICAL ANALYSIS OF CX43 Z-STACKS

A statistical analysis employing both parametric and non-parametric tests was performed to quantitatively assess differences in unique Cx43 puncta count in Z-stack distribution between groups(Fig.2). Analyzed total tracks count for each subset.

**Figure 2.**
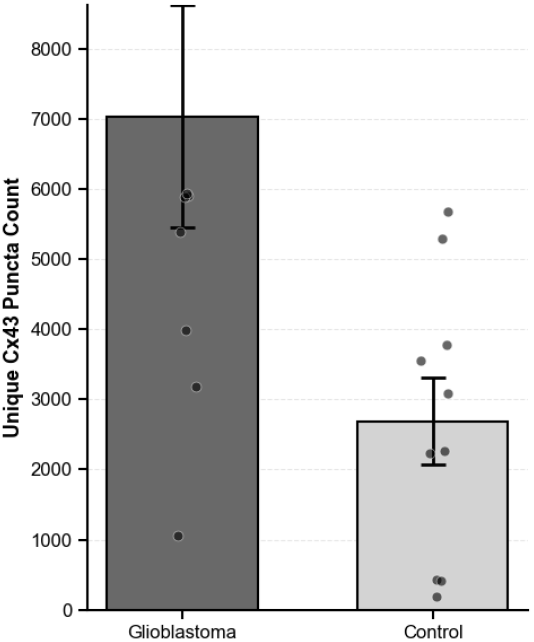
Cx43 puncta count is elevated in glioblastoma compared to control. Bar graph showing mean ± SEM unique Cx43 puncta count in glioblastoma (n=10) and control (n=10) groups. Individual data points are overlaid. Two-tailed independent samples t-test: t = 2.55, p = 0.02, Cohen’s d = 1.14 (large effect size). Error bars represent standard error of the mean (SEM).

Descriptive statistics revealed the following values for pathology (n=10) and control (n=10) groups: mean object counts were 7035.20 ± 5012.17 and 2691.00 ± 1960.82, respectively. Median values were 5889.00 for pathology and 2676.00 for control. Normality assessment using the Shapiro-Wilk test indicated that both samples conformed to normal distribution (pathology: W = 0.8628, p = 0.0824; control: W = 0.9251, p = 0.4017). Levene’s test confirmed homogeneity of variances between groups (F = 1.4368, p = 0.2462). Student’s t-test for independent samples revealed statistically significant differences between groups (t = 2.5525, p = 0.0200). The non-parametric Mann-Whitney U test corroborated these findings (U = 86.0000, p = 0.0073). Effect size, evaluated using Cohen’s d coefficient (d = 1.1415), indicates a large effect magnitude. The mean difference was 4344.20 with a 95% confidence interval [1122.08, 7566.32]. Power analysis demonstrated that the current statistical power was 0.9485 (94.9%), substantially exceeding the recommended threshold of 0.80. The minimum required sample size to achieve 80% power is 9 observations per group.

## 5 CONCLUSION

Our dataset of confocal microscopic images of Cx43 puncta represents a unique resource for neuroscience research, distinguished by its specialized focus and high resolution. The hierarchical organization and detailed documentation enable a wide range of research applications, from basic morphological studies to advanced machine learning model development.

The integration of samples from different species, physiological states, and brain regions provides unprecedented opportunities for comparative studies and the development of sophisticated analytical tools. While not directly compatible with some existing datasets due to methodological differences, our collection complements these resources and opens new avenues for research on astrocytic communication networks.

By making this dataset available to the research community, we aim to accelerate advances in understanding the crucial role of astrocytic gap junctions in normal brain function and pathological conditions, ultimately contributing to the development of new diagnostic tools and therapeutic approaches for neurological disorders.

## ACKNOWLEDGMENTS

This research was funded by the grant of the Ministry of Science and Higher Education of the Russian Federation No. FZNE-2024-0004.

## A APPENDICES

### A.1 Dataset samples and their distribution

### A.2 Existing Astrocyte Datasets

**Table 1.**
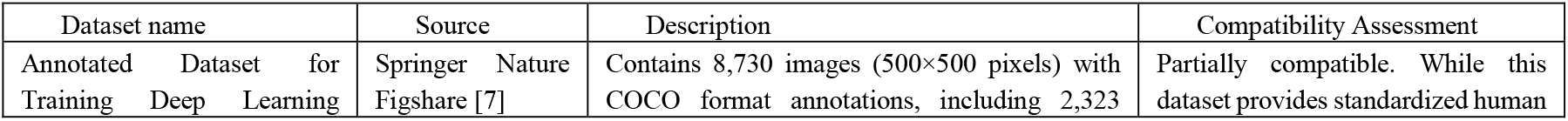
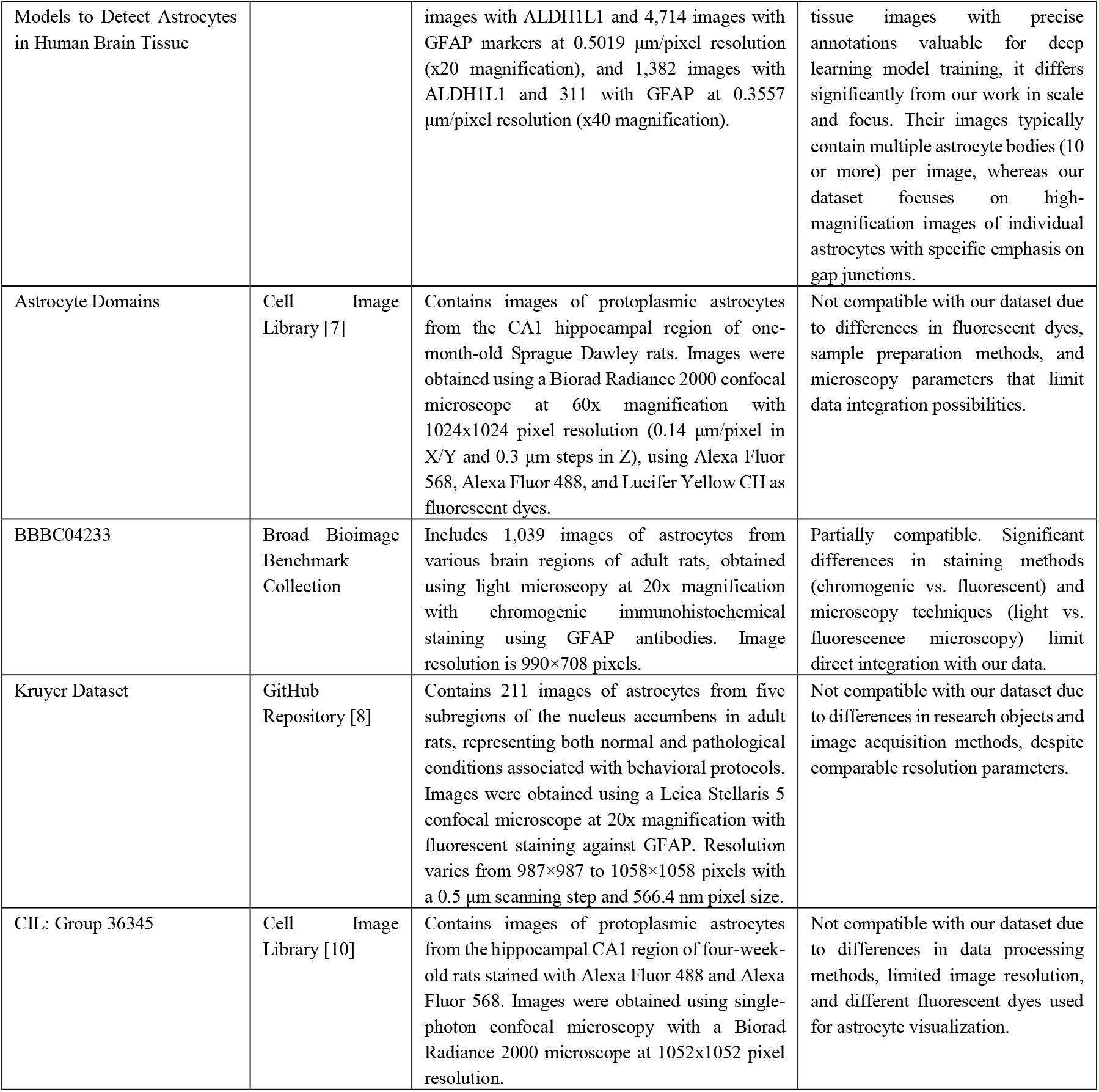
Existing Astrocyte Datasets.

### A.3 Dataset Cataloging Structure

The astrocyte dataset is organized in a strictly hierarchical structure consisting of upper and lower levels, ensuring data systematization, convenient navigation, and subsequent use for analytical and research purposes. The dataset containing 1,058 images of brain astrocytes from animals and humans was created by scanning 20 astrocyte cells.

The structure is organized into five hierarchical levels:

- Species Level (Top): two main categories corresponding to the species from which biological material was collected for astrocyte research using confocal microscopy: human and rat.
- Physiological State Level: subcategories describing the physiological state of the organism at the time of biological material collection:
  - For humans: cancer (anonymized data on people suffering from cancer, focusing on astrocytes in tumor tissues, particularly glioblastoma).
  - For rats: healthy (data from healthy animals used as control samples for research).
- Brain Region Level: data structured by sample localization, specifying the specific brain area from which material was collected for scanning:
  - For humans: glioblastoma (tumor tissues primarily in brain cortex).
  - For rats: brain_cortex.
- Scanning Method Level: indicates the microscopy methods used to obtain astrocyte images:
  - Confocal microscopy: standard microscopy using lasers and optical layers to obtain clear images.
- Fluorescent Marker Level: contains data on fluorescent markers used to stain cells before scanning:
  - Cx43: marker used for labeling connecting proteins such as connexin-43, with orange fluorescent emission (Star Orange)

